## Supplementary Information for "Evidence for absence of links between striatal dopamine synthesis capacity and working memory capacity, spontaneous eye-blink rate, and trait impulsivity"

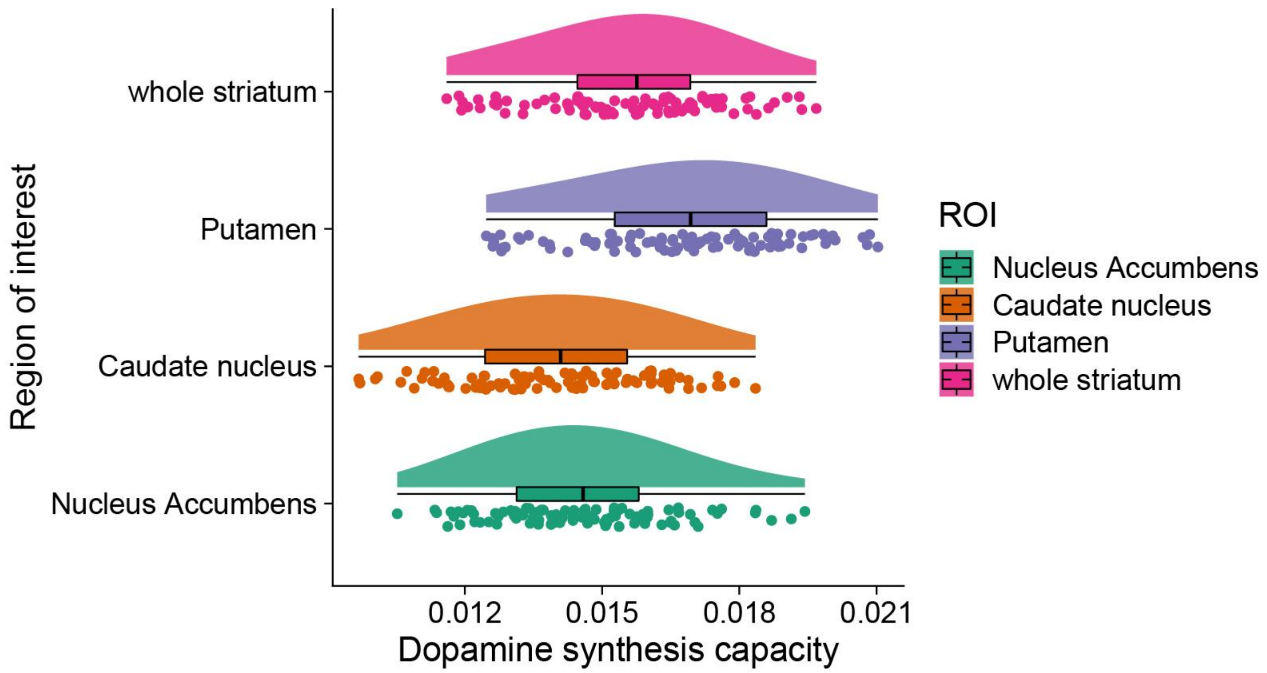


***Supplementary Figure 1.*** *Distribution of dopamine synthesis capacity ($k_i^{cer}$) values in the striatal regions of interest (ROI): caudate nucleus, putamen and nucleus accumbens. The whole striatum ROI is the combination of all three subregions of interest.*

**Table S1. Pearson correlation coefficients for dopamine synthesis capacity in the three striatal regions of interest.**

| | Caudate nucleus | Putamen | Nucleus Accumbens |
|---|---|---|---|
| Caudate nucleus | 1 | 0.751 | 0.649 |
| Putamen | 0.751 | 1 | 0.787 |
| Nucleus Accumbens | 0.649 | 0.787 | 1 |

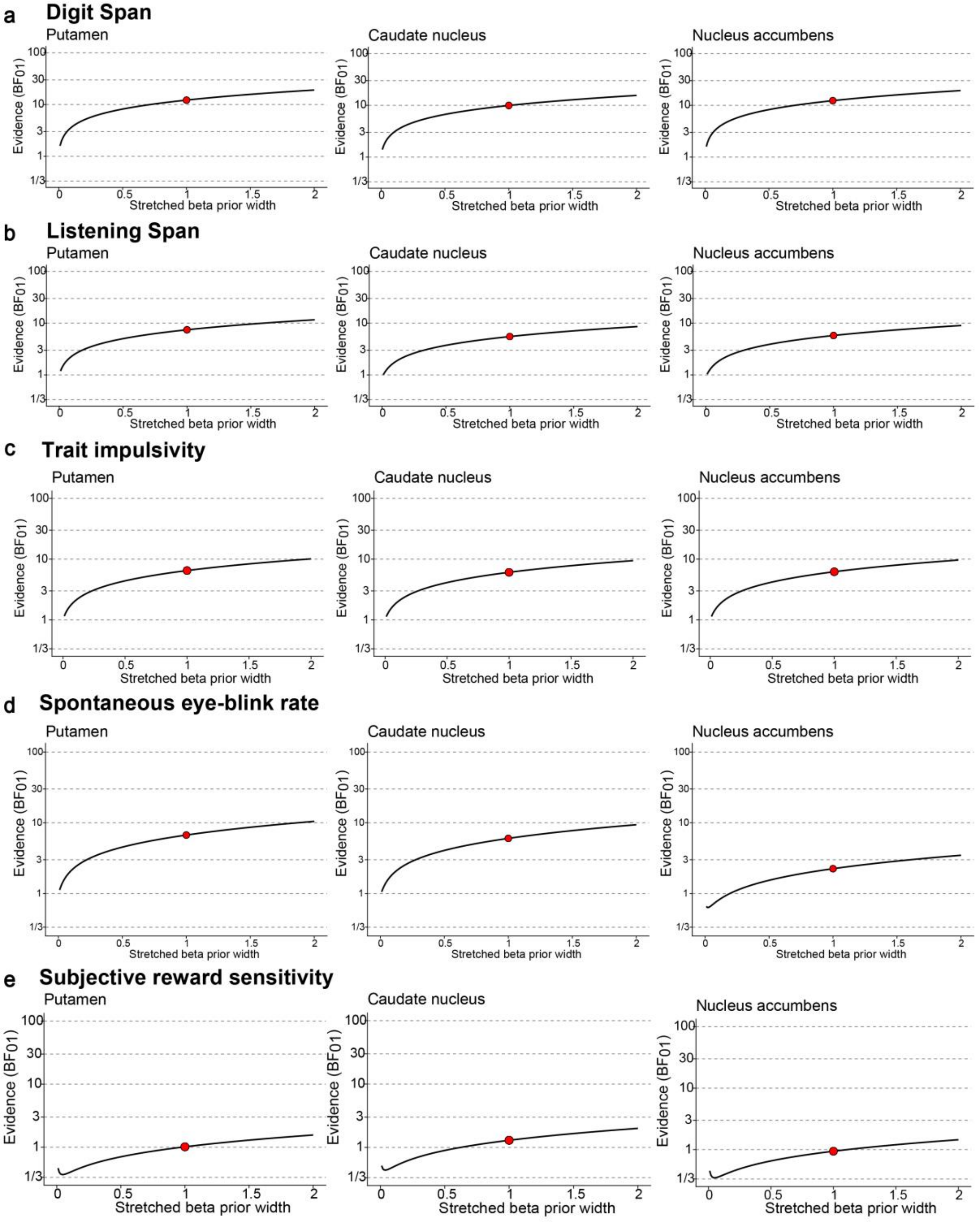


***Supplementary Figure 2. Bayes Factor robustness checks.*** *Bayes Factor results as function of varying priors for the null hypothesis of no positive correlation (H0) versus a positive correlation (H1) between striatal dopamine synthesis capacity and **(a)** working memory capacity measured with the Digit Span task (N = 94), **(b)** working memory capacity measured with the Listening Span task (N = 94), **(c)** trait impulsivity measured with the BIS-11 questionnaire (N = 66), **(d)** spontaneous eye-blink rate (N = 92), or **(e)** subjective reward sensitivity measured with the Behavioural Activation Scale (N = 94). The red dot corresponds to the Bayes Factor for the default prior specification.*

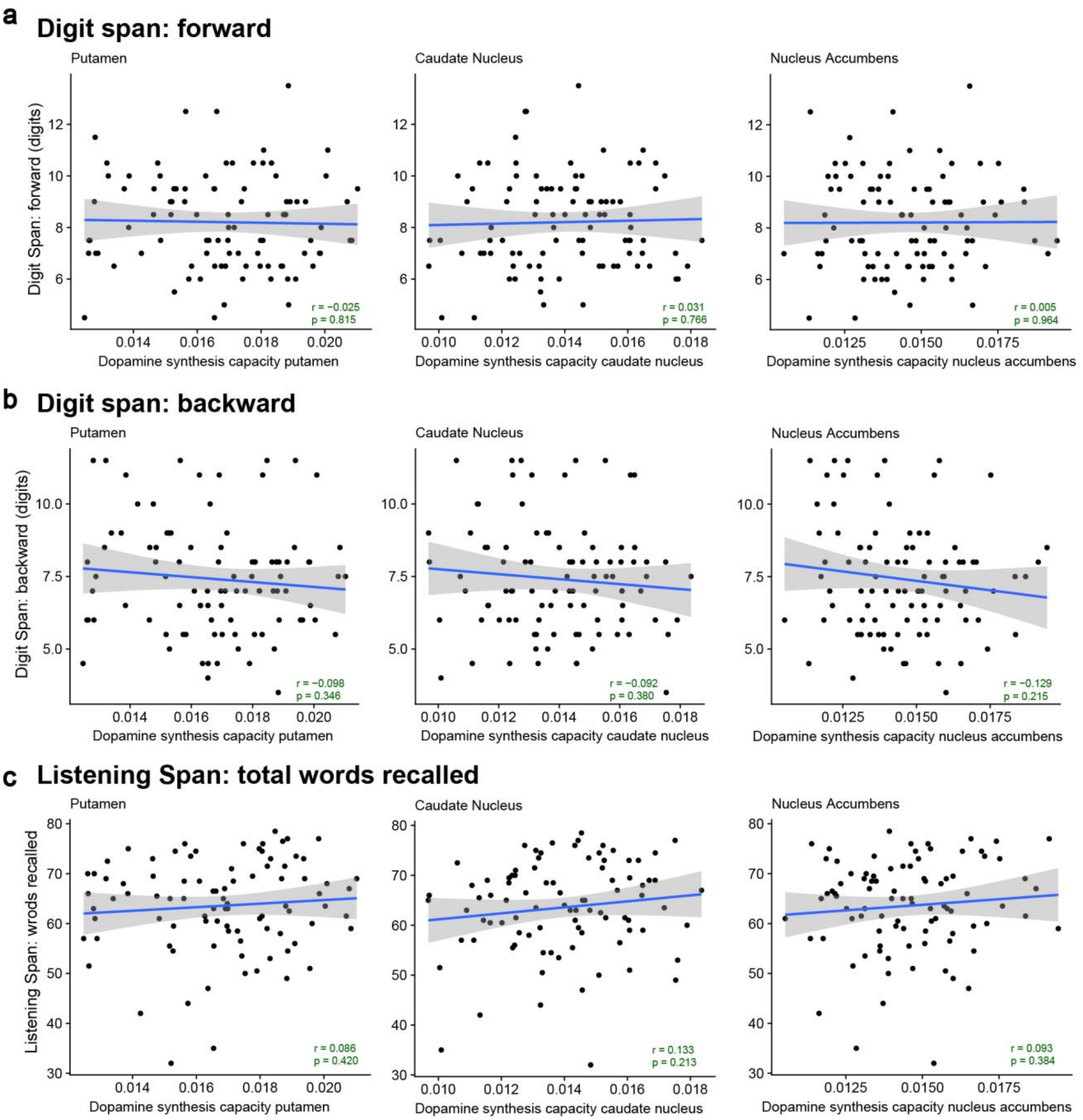


***Supplementary Figure 3. No significant correlations between striatal dopamine synthesis capacity and subsections of the working memory tasks (N=94).*** *Correlations between dopamine synthesis capacity in the putamen, caudate nucleus or nucleus accumbens ROIs and* ***(a)*** *the forward section and* ***(b)*** *backward section of the Digit Span task and* ***(c)*** *the total number of words recalled in the Listening Span task. The p-values of the two-sided Pearson correlation coefficients are not corrected for multiple comparisons.*

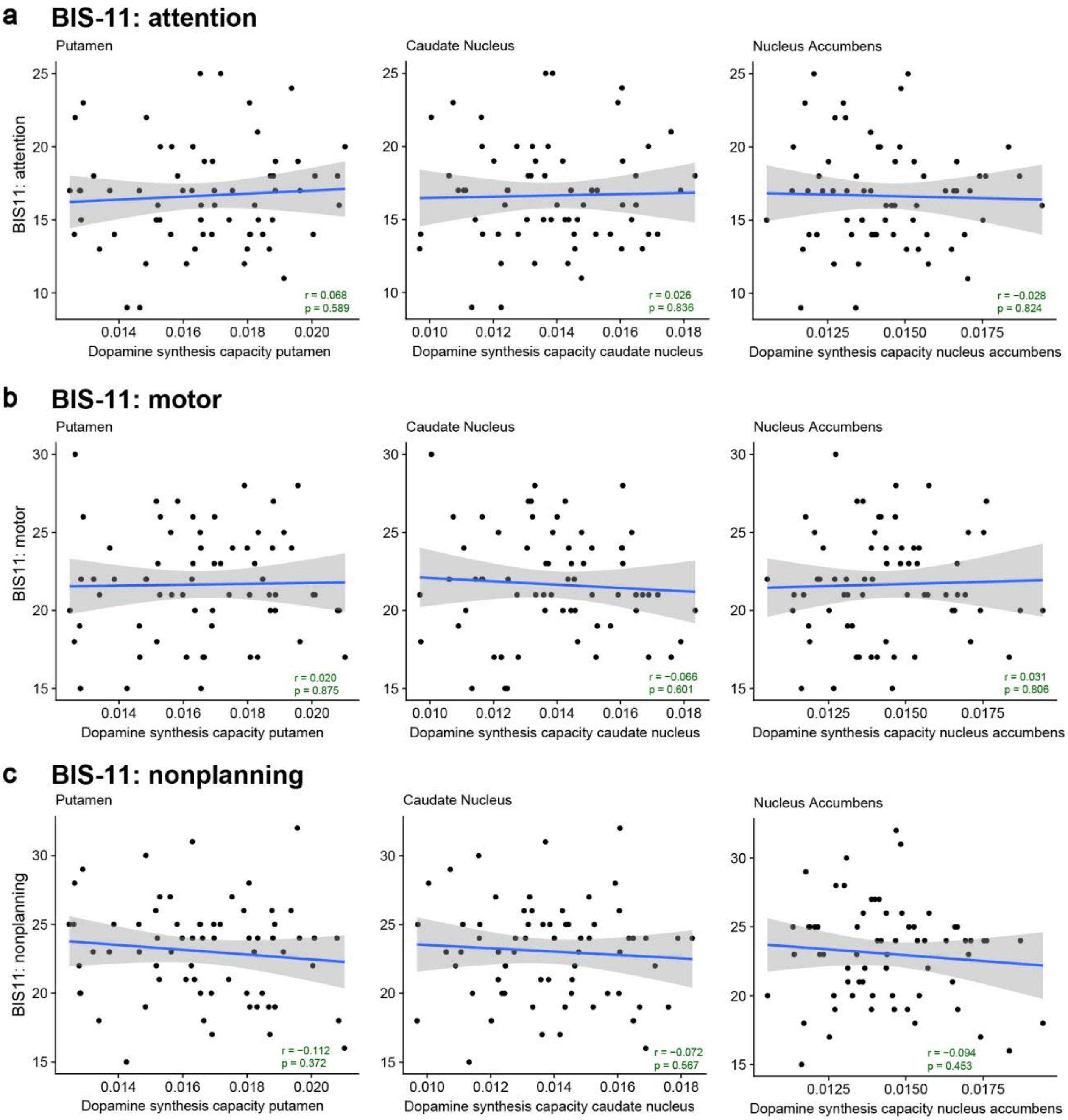


***Supplementary Figure 4. No significant correlations between striatal dopamine synthesis capacity and subsections of the BIS-11 questionnaire of trait impulsivity (N=66).*** *Correlations between dopamine synthesis capacity in the putamen, caudate nucleus or nucleus accumbens ROIs and the BIS-11 sub-scale on* ***(a)*** *attention,* ***(b)*** *motor actions, and* ***(c)*** *nonplanning. The p-values of the two-sided Pearson correlation coefficients are not corrected for multiple comparisons.*

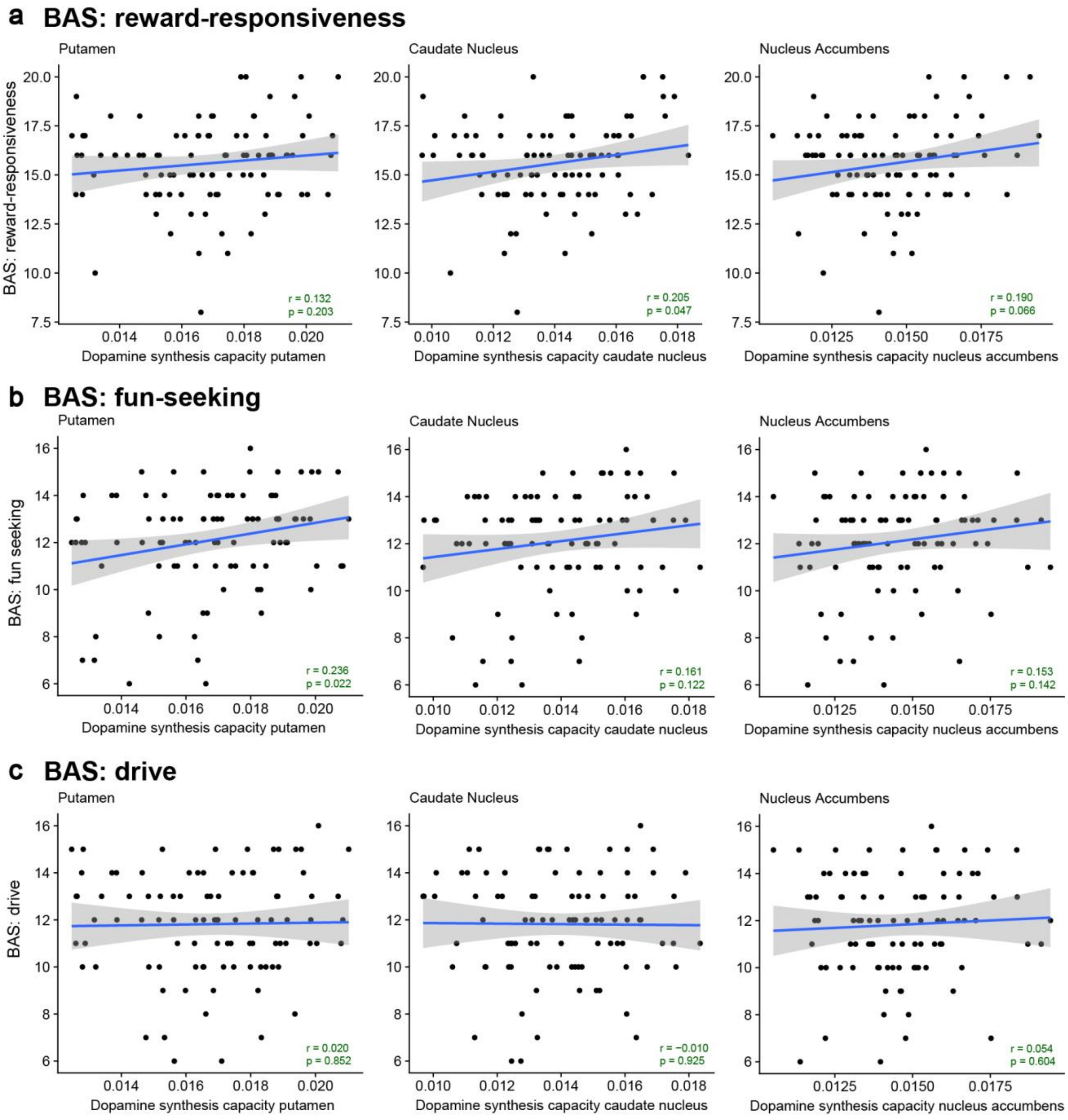


***Supplementary Figure 5. No significant correlations between striatal dopamine synthesis capacity and subsections of the BAS questionnaire of trait reward sensitivity (N=94).*** *Correlations between dopamine synthesis capacity in the putamen, caudate nucleus or nucleus accumbens ROIs and the BAS sub-scale on* ***(a)*** *reward-responsiveness,* ***(b)*** *fun-seeking, and* ***(c)*** *drive. The p-values of the two-sided Pearson correlation coefficients are not corrected for multiple comparisons.*

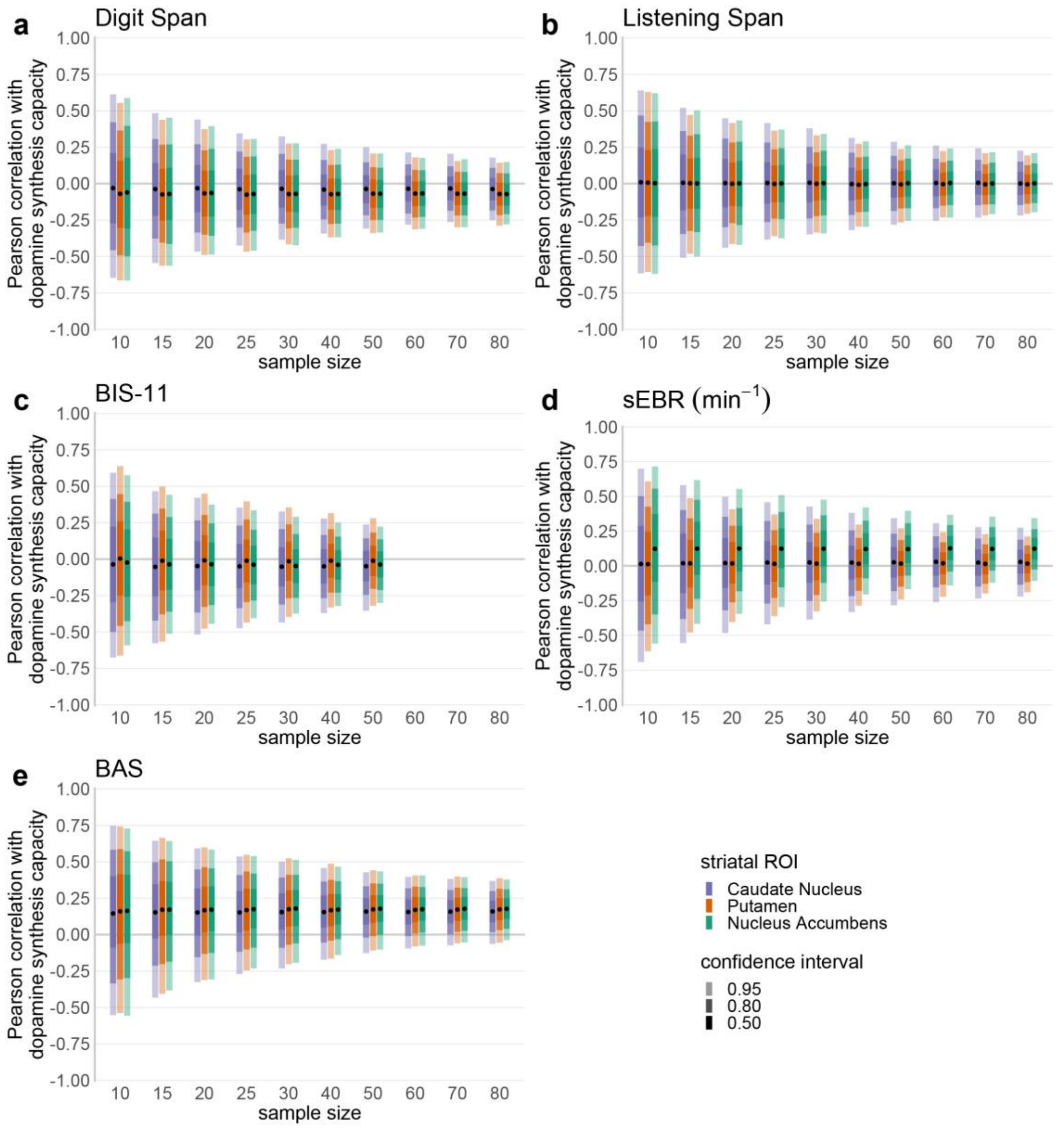


***Supplementary Figure 6. Sampling variability of the correlation between dopamine synthesis and trait measures.*** *From the full dataset, random subsamples were drawn for each of a series of sample sizes (N=10, 15, 20, 25, 30, 40, 50, 60, 70, and 80), and for each subsample the correlations between the trait measures and dopamine synthesis capacity was computed. This was repeated 5000 times for each sample size to obtain distributions of the correlations between the trait measures and dopamine synthesis capacity. These distributions were then summarized with the 95%, 80%, and 50% confidence intervals to reveal the sampling variability of the correlations at varying sample sizes. This analysis was performed for each of the key trait measures under consideration:* ***(a)*** *working memory capacity measured with the Digit Span task,* ***(b)*** *working memory capacity measured with the Listening Span task,* ***(c)*** *trait impulsivity measured with the BIS-11 questionnaire,* ***(d)*** *spontaneous eye-blink rate, and* ***(e)*** *subjective reward sensitivity measured with the Behavioral Activation Scale. As indicated in the bottom-right panel, the different colors represent the striatal region of interest (ROI) in which dopamine synthesis capacity was estimated, and the opacity represents the confidence interval. The black dots represent the means across subsamples. Note that for BIS-11, the subsampling analysis was performed for a reduced range of sample sizes, because the original data for this measure was limited to N=66 (see Methods for details).*

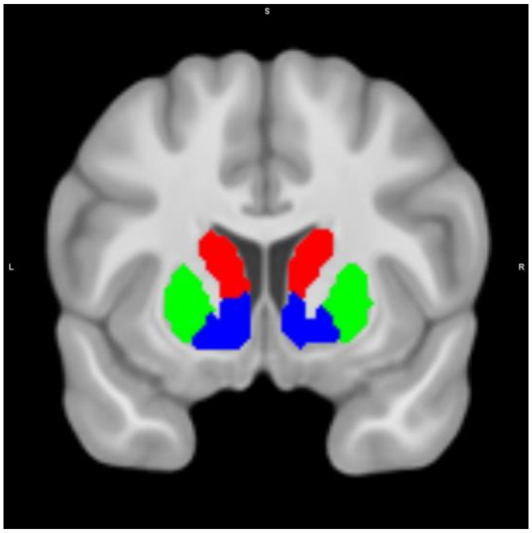


***Supplementary Figure 7. Analysis masks of the striatal regions of interest: caudate nucleus (red), putamen (green), and nucleus accumbens (ventral striatum; blue).*** *The masks are based on an independent, functional connectivity-based parcellation of the striatum (Piray et al., 2017).*
